# Prolonged exposure to hypergravity increases biomass and alters biomass allocation in *Arabidopsis thaliana* (L.) Heynh. with no apparent impact on element content in the shoot system

**DOI:** 10.1101/2024.06.18.599652

**Authors:** Kazuki Ohara, Mizuki Katayama, Hiroyuki Kamachi, Atsushi Kume, Ichirou Karahara

## Abstract

Previous studies have already shown that plants can complete their life cycle under microgravity. However, the effects of long-term exposure to altered gravity conditions, including microgravity, on most of the biological processes of a plant’s life cycle remain largely unexplored. Given the limited opportunities for space experiments, it is crucial to conduct ground-based experiments, such as hypergravity experiments. To investigate the longer-term effects of hypergravity, we have developed and utilized a custom-built hypergravity cultivation system using a centrifuge equipped with lighting, enabling the continuous growth of seed plants under hypergravity conditions. In this study, we examined the effects of 10 *g* hypergravity on the biomass of the shoot system (stems and rosette leaves) and the root system of vascular plants for the first time, covering the entire cultivation period from germination to 40 days. Our results showed that the dry mass of the stem per unit length was significantly higher under 10 *g* compared to the 1 *g* control, indicating a typical gravity resistance response of the stem. Moreover, the total dry mass of the stems, rosette leaves, and roots was higher under 10 *g* hypergravity compared to the 1 *g* control, suggesting an increase in biomass at the individual plant level. We also observed that the leaf mass per area of the rosette leaf was higher under hypergravity compared to the 1 *g* control, indicating enhanced photosynthesis rates in Arabidopsis and resulting in increased biomass of individual plants. In terms of biomass allocation, both root-shoot ratio and root mass fraction were significantly higher under hypergravity conditions compared to the 1 *g* control. Furthermore, we measured the content of mineral elements (Ca, Co, Cu, Fe, K, Mg, Mn, Mo, P, Zn) in the roots and rosette leaves using inductively coupled plasma optical emission spectrometry. Despite the increase in dry mass of the root system, we found no significant differences in the content of any of the ions analyzed between 10 *g* and 1 *g* conditions, indicating that mineral nutrient uptake homeostasis is maintained even under hypergravity conditions.

## 1. Introduction

Cultivating plants in space habitats is crucial for the development of bioregenerative life support systems necessary for long-term space exploration and utilization. For a considerable time, there was uncertainty about whether plants accustomed to Earth’s gravitational acceleration (1 *g*) could complete their life cycles under altered gravitational conditions. However, through successive space experiments, including seed-to-seed experiments with plant species like Arabidopsis (Merkys and Laurinavicius, 1983; Link et al., 2003; Musgrave and Kuang, 2003; Link et al., 2014) and *Brassica rapa* L. (Musgrave et al., 2000; Musgrave et al., 2005), it has been shown that Earth’s gravity is not crucial for plant growth and reproduction. Nonetheless, the precise effects of altered gravity on different morphological and physiological aspects of plant development are still largely unknown. Therefore, it is crucial to examine these effects to better understand and maximize plants’ adaptive capability to altered gravitational environments for future manned space exploration endeavors.

Given the limited opportunities for space experiments, conducting ground-based experiments is vital. Hypergravity experiments offer the only method to simulate altered gravitational acceleration conditions over prolonged periods for cultivating plants on Earth. Allen et al. (2009) examined effects of 4 *g* hypergravity for up to 16 days (d) on the growth of *Brassica rapa* and *Arabidopsis thaliana* plants employing a large-diameter centrifuge available at a space center. However, the availability of such facilities is limited. Therefore, to perform lab-based prolonged hypergravity experiment we first used a commercial centrifuge attached with a lighting system, demonstrated the effect of exposure to 8 *g* for 10 d on lignin deposition in the peduncle, and on tissue anatomy (Shinohara et al., 2024). We also developed a custom-built centrifuge equipped with a lighting system and demonstrated that rhizoid lengths of *Physcomitrium* (*Physcomitrella*) *patens* (Hedw.) Bruch et Schimp. were significantly increased under 10 g and the area-based photosynthesis rate was also enhanced under 10 *g* (Takemura et al., 2017; Takemura et al., 2017). However, the first custom-built centrifuge equipped with a lighting system, MIJ-7, has a slight problem of vibration generated during centrifugation (Mori et al., 2017). Therefore, the authors have further developed custom-built centrifuges having lighting systems with only negligible vibration, MK-2 and MK-3, which is more suitable for longer period of hypergravity cultivation with seed plants (Mori et al., 2017; Takemura et al., 2017).

Understanding the effects of altered gravity on plant biomass and crop productivity is crucial for space agriculture and human space exploration. In this study, we investigated the effects of 10 *g* hypergravity on the biomass of the shoot system (stems and rosette leaves) and the root system of seed plants from germination to a cultivation period of up to 40 d using our improved hypergravity cultivation system, MK-2. To understand the physiological status of plants grown under specific environments, biomass allocation has been a focus from an ecological perspective (Ericsson, 1995; ÅGren and Franklin, 2003; Mašková and Herben, 2018), although it has not been explored in the context of gravitational environments. Biomass allocation indicators such as root-shoot ratio and leaf dry mass per area (LMA) are commonly analyzed for this purpose. The root-shoot ratio and its inverse, the shoot-root ratio, are frequently used to assess biomass allocation in a whole plant (Poorter et al., 2012), which can be influenced by various environmental factors including water deficit, nutrient availability, light, and CO_2_ levels (Wilson, 1988). More detailed indicators such as stem mass fraction (SMF), leaf mass fraction (LMF), and root mass fraction (RMF) are also employed to evaluate biomass allocation (Poorter and Nagel, 2000). LMA, an important functional indicator, is widely used in plant ecology, agronomy, and forestry, responds to environmental factors such as light, CO_2_ levels, and water availability (Poorter et al., 2009). These indicators were examined to understand the physiological status of plants grown under prolonged hypergravity conditions and to correlate them with other environmental conditions.

Additionally, understanding the effects of altered gravitational conditions on mineral nutrient acquisition is crucial for the development of space agriculture. This aspect has not been previously investigated, despite its significance in biomass accumulation, as nutrient acquisition directly impacts plant growth (Theocharis and Ioannis, 2013). Moreover, considering the cultivation of plants using moon regolith, it is important to assess the effects of altered gravity on mineral nutrient acquisition due to potential metal stress resulting from the uptake of heavy metals from the regolith (Paul et al., 2022). Therefore, we investigated whether the content of inorganic elements in the shoot is affected under hypergravity conditions using an ionomics approach with an Inductively Coupled Plasma Optical Emission Spectroscopy (ICP-OES) instrument for the first time.

## 2. Materials and Methods

### 2-1. Plant materials and hypergravity treatment

Seeds of Arabidopsis (*Arabidopsis thaliana* (L.) Heynh.) of Columbia-0 line were surface sterilized with 99 % (v/v) ethanol for 10 s. Four seeds were planted on 30 mL of 1.0 % (w/v) agar containing Murashige and Skoog medium (Fujifilm Wako Pure Chemical, Osaka, Japan) in a polycarbonate container (Agripot, Kirin Beer Co., Ltd, Tokyo, Japan; 70 mm in diameter, 120 mm in height), kept for 3 d at 4 °C in the dark, and then allowed to grow at 25 °C for 30 or 40 d under hypergravity at 10 *g* in the continuous light (F32W-T fluorescent bulb, Nichido Ind. Co. Ltd., Osaka, Japan) with the intensity being 120 µmol m^−2^ s^−1^ at plant level using a centrifugal plant cultivation system, MK2 (Matsukura Co., Ltd., Kurobe, Japan). Because the ventilation hole at the top a container receives a wind at a speed of 2.96 m s^−1^ during centrifugation for hypergravity conditions at 10 *g*, a wind at the speed of 2.9 - 3.0 m s^−1^ is applied for the 1 *g* control using an air circulator (530-JP, Vornado Air LLC, Andover, USA) (Fig. 1a). The ventilation hole was covered with a cotton ball (10 mm in diameter,) and sealed with surgical tape (Micropore Surgical Tape, 3M, ) to prevent contamination as well as excess ventilation.

**Fig. 1.**
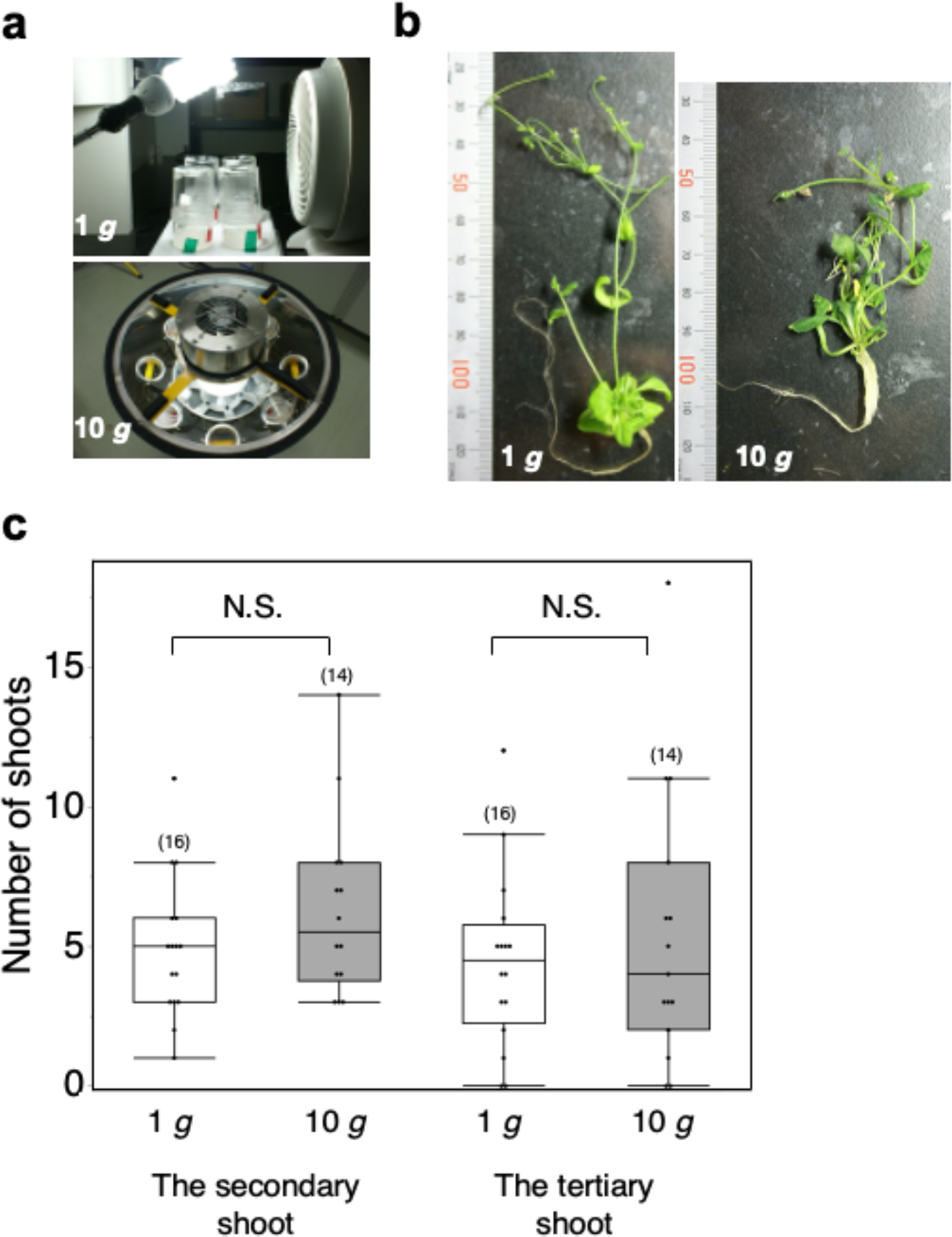
Setup of the hypergravity experiment (**a**) and the morphology of a typical 40-d-old Arabidopsis plant under 1 *g* control conditions or following exposure to 10 *g* hypergravity (**b**). (**c**) Effects of 40-d exposure to 10 *g* hypergravity on the numbers of the secondary shoots and the tertiary shoots. N.S., *P* > 0.05, Wilcoxon rank sum test. Each box plot shows 1.5 interquartile range of the data points (whiskers), third quartile, median, first quartile value, and an outlier. Dots show individual data points. The sample sizes are indicated in parentheses.

### 2-2. Measurements of morphology and dry mass of primary inflorescence main stems, rosette leaves, and root system

The main axis of the primary inflorescence, which comprises the rachis and peduncle, is referred to as the main stem for simplicity and understandability. After the hypergravity treatment, the lengths of the main stem of the primary inflorescence and the longest root were measured, the number of rosette leaves was counted, and the width, length, and area of the rosette leaves were measured. Then, the samples were dried at 60 °C for 24 hours, and the dry mass of the main stem of the primary inflorescence, the entirety of the rosette leaves, and the entirety of the root systems were measured using an ultramicrobalance (SE2, Sartorius AG, Goettingen, FRG). Since detailed morphological analysis of the rosette leaves was not conducted with the 40-d-old plants, only data from the 30-d-old plants are presented.

### 2-3. Measurements of the content of elements in the main stem of the primary inflorescence and the rosette leaves

After measuring the morphology and dry mass of the primary inflorescence main stems, rosette leaves, and root system, the dried main stem of the primary inflorescence and rosette leaves were digested using wet ashing method. Thirty mg of dried samples were placed into a test tube, and 2 mL of concentrated nitric acid were added. The sample was heated to 75 °C and gently mixed until dissolved. Then, 0.3 mL of perchloric acid were added to the sample, and it was heated to 120 °C. The sample solution was gently mixed until it became colorless and transparent. The sample solution was cooled, and 0.1 N nitric acid was added to adjust the volume to 10 mL. The content of elements (Ca, Co, Cu, Fe, K, Mg, Mn, Mo, P, Zn) in the sample was then analyzed using the ICP-OES instrument (Optima 7300DV, Perkin Elmer Instruments Co., Ltd., Yokohama, Japan). The experiment was repeated twice to test reproducibility.

### 2-4. Statistical analysis

For testing statistical difference between two samples having data of continuous variables, normality and variance equality of population distribution was examined. When both of the assumptions held, Student’s *t*-test was performed. When neither of the assumptions held, Wilcoxon rank sum test was performed. When normality held but variance equality did not, as tested by using Bartlett test, Welch’s *t*-test was performed instead. For testing statistical difference between two samples having data of discrete variables, Wilcoxon rank sum test was performed. Box plot shows 1.5 interquartile range of the data points (whiskers), third quartile, median, first quartile value, and an outlier. Statistical analyses were conducted using JMP Pro version 17 software (SAS Institute Inc., Cary, NC, USA).

## 3. Results

### 3-1. Effects of prolonged hypergravity exposure on the growth of the primary inflorescence

The effect of prolonged hypergravity exposure over 40 d on the morphology of the primary inflorescence was assessed. Because lateral shoots were already observed in the plants under 1 *g* control conditions or after 40-d of exposure to 10 *g* hypergravity (Fig. 1b), we investigated whether hypergravity affected branch formation. However, no significant difference was observed in the numbers of secondary or tertiary shoots between the 1 *g* and 10 *g* conditions (Fig. 1c). Similar results were obtained for the 30-d-old plants exposed to 10 *g* for 30 d (Supplementary Fig. 1a).

In terms of the effects of prolonged hypergravity exposure on stem growth, the length of the main stem of the primary inflorescence was significantly shorter under hypergravity conditions compared to the 1 *g* control for 40-d-old plants (Fig. 2a). Although not significant for 30-d-old plants, there was a tendency for the main stem to be shorter under hypergravity (Supplementary Fig. 1b). While the length was significantly shorter only in the 3rd internode under hypergravity compared to the 1 *g* control, it showed a similar trend in other examined internodes (Fig. 2b). Conversely, the total dry mass of the stem was significantly greater under hypergravity compared to the 1 *g* control (Fig. 2c, Supplementary Fig. 1c). Consequently, the dry mass of the stem per unit length was significantly higher under hypergravity compared to the 1 *g* control (Fig. 2d, Supplementary Fig. 1d). For 40-d-old plants, the dry mass of the stem per unit length, including stems of the secondary and tertiary shoots, was examined (Fig. 2a), while for 30-d-old plants, the dry mass of the main stem per unit length was analyzed (Supplementary Fig. 1d). The former value was lower than the latter because it represents an average that includes the secondary and tertiary shoots, possibly indicating that these shoots are thinner than the primary shoot, while the latter only includes the main stem. Additionally, the total dry mass of the stems of the lateral shoots was significantly higher under hypergravity compared to the 1 *g* control when analyzed using 30-d-old plants (Supplementary Fig. 1e), despite no significant difference in the numbers of secondary or tertiary shoots between the 1 *g* and 10 *g* conditions (Supplementary Fig. 1a).

**Fig. 2.**
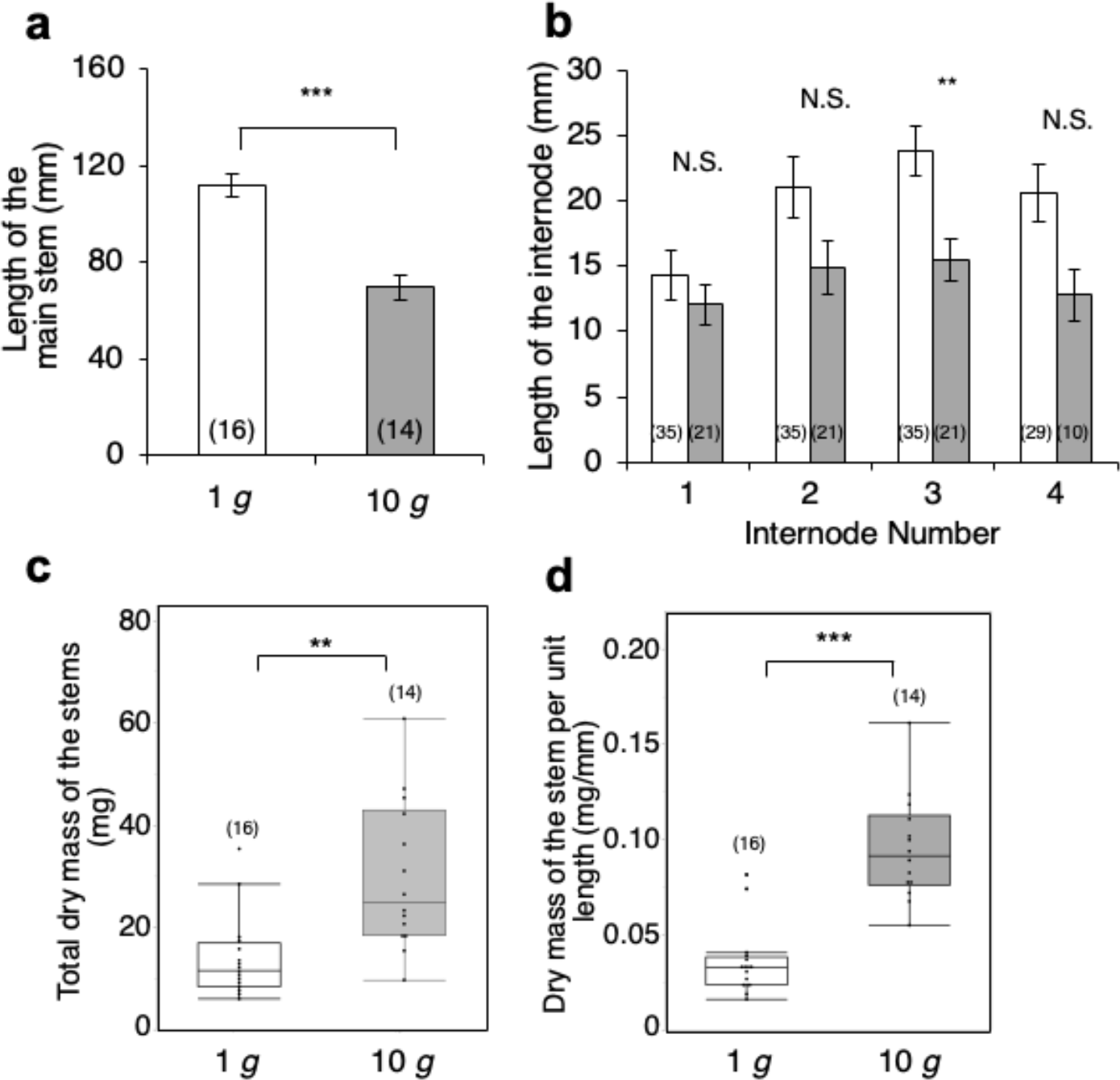
Effects of 40-d exposure to 10 *g* hypergravity on the main stem of the Arabidopsis primary inflorescence. The sample sizes are indicated in parentheses. (**a**) The length of the main stem of the primary inflorescence. ***, *P* < 0.001, Student’s *t*-test. (**b**) The internode lengths of the main stem. **, *P* < 0.01, Student’s *t*-test. N.S., *P* > 0.05, Wilcoxon rank sum test. (**c**) The total dry mass of the main stem. ***, *P* < 0.001, Wilcoxon rank sum test. (**d**) The dry mass of the main stem per unit length. ***, *P* < 0.001, Wilcoxon rank sum test. (**a, b**) Values are presented as mean ± SE.

Regarding the effects of prolonged hypergravity exposure on the rosette leaf, there was no significant difference in the number of rosette leaves between the 1 *g* and 10 *g* conditions (Fig. 3b), while the total dry mass of the rosette leaves was significantly higher under hypergravity compared to the 1 *g* control (Fig. 3c, Supplementary Fig. 1f). Because the shape of the rosette leaf was only analyzed using 30-d-old plants, data on the area, length, and width of the rosette leaves from these plants were included in Fig. 3. The length of the rosette leaf was significantly greater under hypergravity compared to the 1 *g* control (Fig. 3e), while there was no significant difference in the width between the 1 *g* and 10 *g* conditions (Fig. 3f). Consequently, the area of the rosette leaf was significantly larger under hypergravity compared to the 1 *g* control (Fig. 3d), and the LMA of the rosette leaf was significantly higher under hypergravity compared to the 1 *g* control (Fig. 3g).

**Fig. 3.**
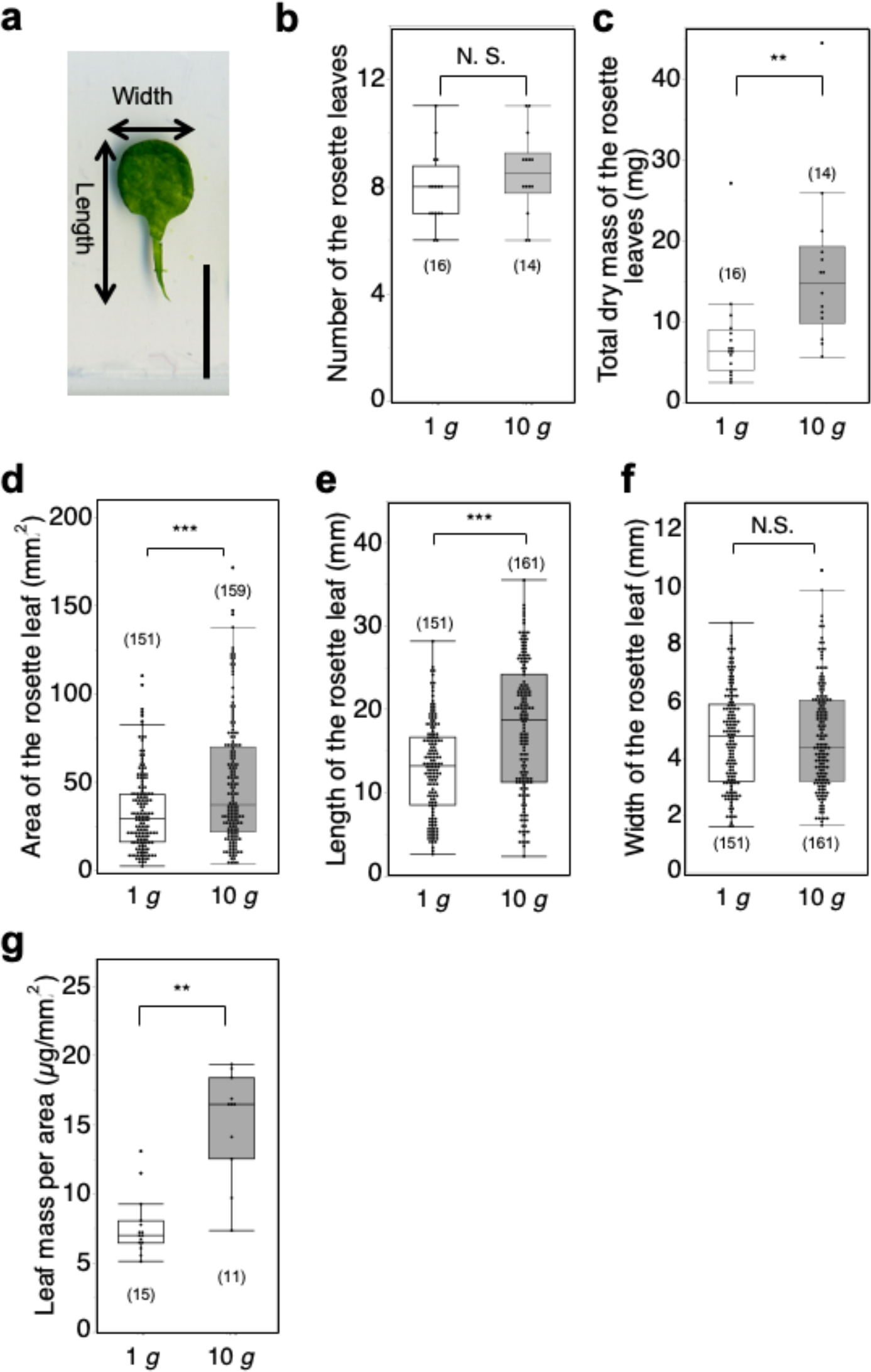
Effects of 30- or 40-d exposure to 10 *g* hypergravity on the rosette leaves of the Arabidopsis plants. The sample sizes are indicated in parentheses. (**a**) The width and length measured for the rosette leaves. Scale bar 10 mm. (**b**) Number of the rosette leaves (40-d exposure). (**c**) The total dry mass of the rosette leaves (40-d exposure). (**d**) The area of the rosette leaf (30-d exposure). (**e**) The length of the rosette leaf (30-d exposure). (**f**) The width of the rosette leaf (30-d exposure). (**g**) The leaf dry mass per area (30-d exposure). N.S., *P* > 0.05; **, *P* < 0.01; ***; *P* < 0.001; Wilcoxon rank sum test.

We collected data on the dry mass of the entirety of the cauline leaves only from the 30-d-old plants. However, due to the lack of consistent results regarding the effects of hypergravity, we did not present these data and excluded them from the total biomass comparison with the 40-d-old plants. Nevertheless, using the same experimental dataset as Supplementary Fig. 1f, we calculated mean ratios of the dry mass of the cauline leaves to that of the rosette leaves, which were 0.48 ± 0.07 (mean ± SE, n = 15) for the 1 *g* condition and 0.19 ± 0.06 (mean ± SE, n = 8) for the 10 *g* condition, for reference.

The effects of prolonged hypergravity exposure on root system growth were examined. Due to challenges in completely retrieving fine roots from the agar media, the accuracy of the root data is limited. However, given the consistent difficulty experienced across gravitational conditions, exploring the effects on the root system remains valuable. As a result, no significant difference was found in the length of the longest root between 1 *g* and 10 *g* conditions when examined using 40-d-old plants (Fig. 4a), while it was higher under 10 *g* conditions when examined using 30-d-old plants (Supplementary Fig. 1d). Despite this inconsistency, the dry mass of the entire root system was significantly higher under hypergravity conditions compared to the 1 *g* control, both in the 40-d-old and 30-d-old plants (Fig. 4b, Supplementary Fig. 1h).

**Fig. 4.**
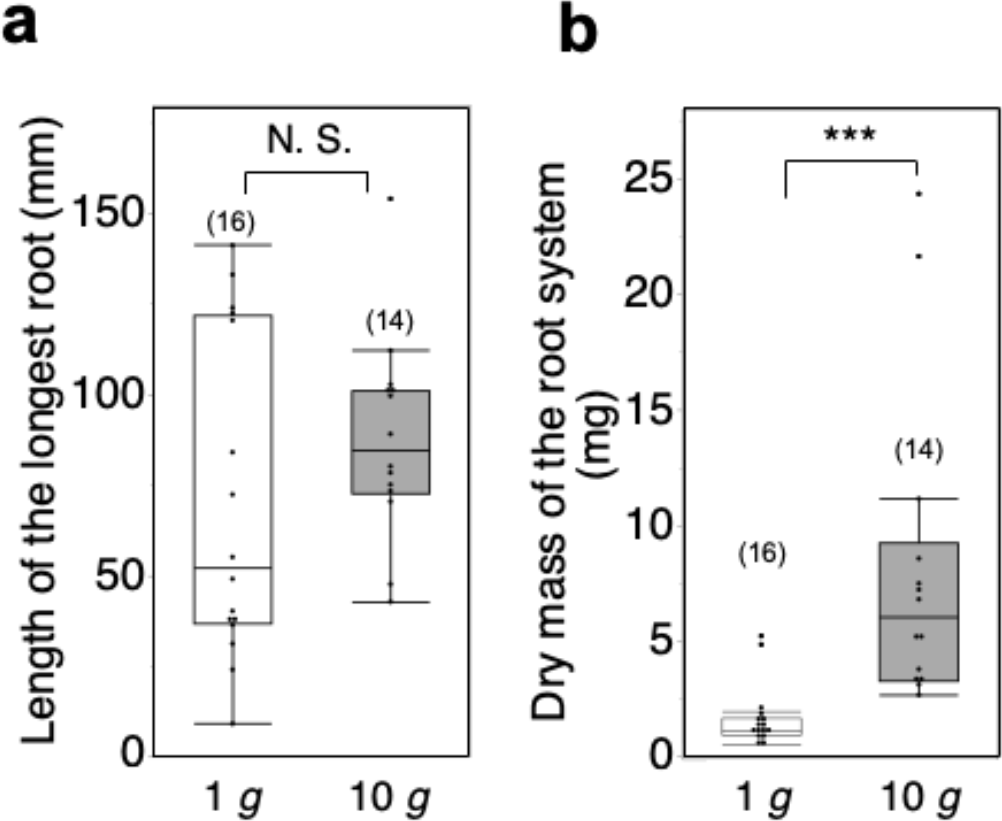
Effects of 40-d exposure to 10 *g* hypergravity on the root system of the Arabidopsis plants. (**a**) The length of the longest root. (**b**) The dry mass of the entirety of the root system. The sample sizes are indicated in parentheses. N.S., *P* > 0.05; ***; *P* < 0.001; Wilcoxon rank sum test.

### 3-2. Effects of prolonged hypergravity exposure on biomass allocation

As mentioned above, the dry mass of any of the stems, rosette leaves, or roots was significantly higher under hypergravity conditions compared to the 1 *g* control. As a natural consequence, the total dry mass of the entirety of the stems, roots, and rosette leaves, which is comparable to the biomass of an individual plant, was significantly higher under hypergravity compared to the 1 *g* control (Fig. 5a, Supplementary Fig. 2a). In order to determine whether biomass allocation is affected under hypergravity conditions, root-shoot ratio, SMF, LMF, and RMF were calculated. As a result, root-shoot ratio was significantly higher under hypergravity conditions compared to the 1 *g* control (Fig. 5b, Supplementary Fig. 2b). For 30-d-old plants, SMF was significantly lower and LMF was significantly higher under hypergravity conditions compared to the 1 *g* control (Supplementary Fig. 2c, d). However, these significant differences for both factors disappeared in the case of 40-d-old plants (Fig. 5c, d). Despite this, RMF was significantly higher under hypergravity conditions compared to the 1 *g* control in both 30-d-old and 40-d-old plants (Fig. 5e, Supplementary Fig. 2e).

**Fig. 5.**
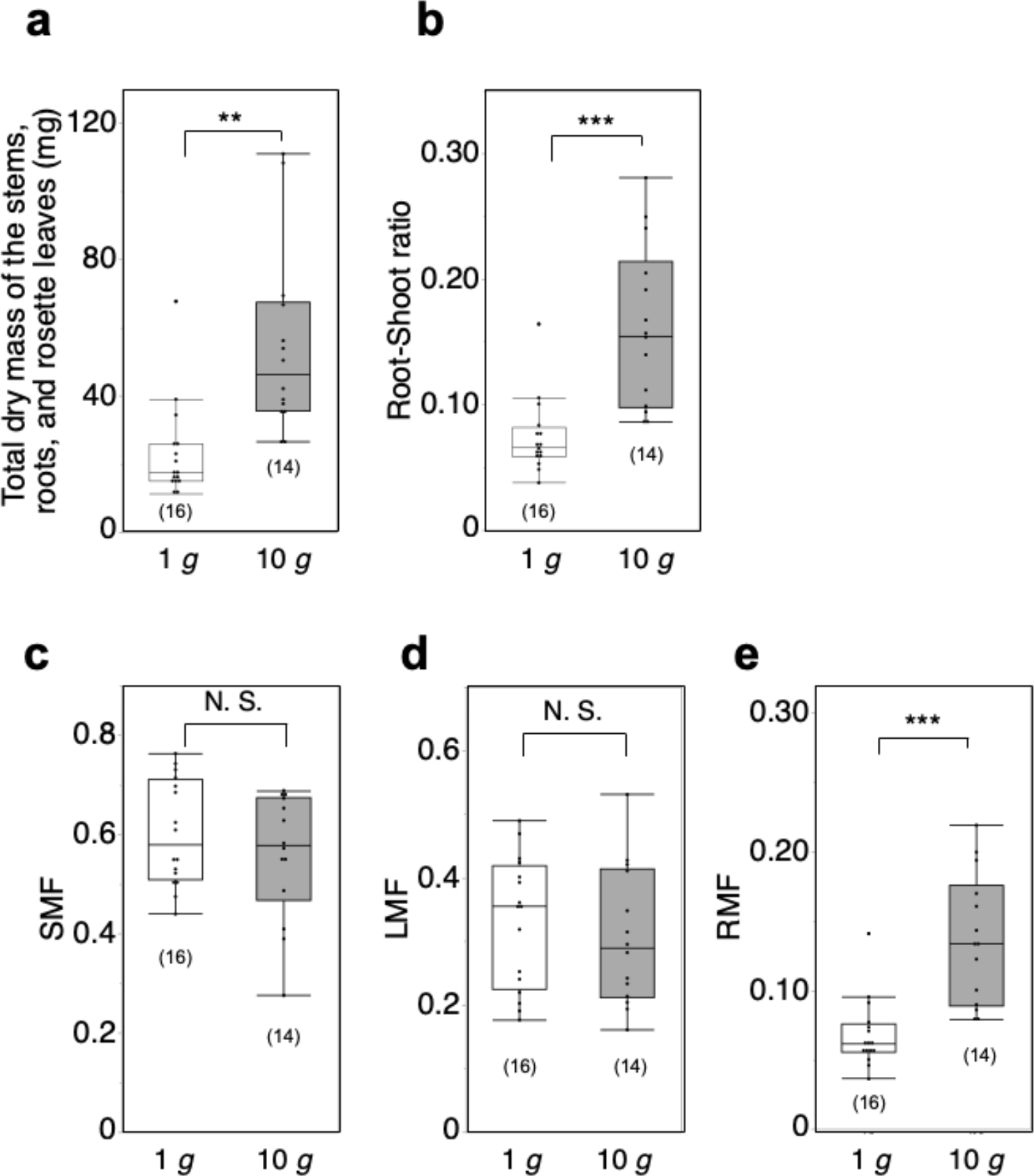
Effects of 40-d exposure to 10 *g* hypergravity on the biomass allocation of the Arabidopsis plants. (**a**) The total dry mass of the stems, roots, and rosette leaves. **; *P* < 0.01, Wilcoxon rank sum test. (**b**) Root-Shoot ratio. ***, *P* < 0.001, Student’s *t*-test of angler transformed values of the data. (**c**) stem mass fraction (SMF). N.S., *P* > 0.05, Student’s *t*-test of angler transformed values of the data. (**d**) leaf mass fraction (LMF) N.S., *P* > 0.05, Student’s *t*-test of angler transformed values of the data. (**e**) root mass fraction (RMF). ***, *P* < 0.001, Welch’s *t*-test of angler transformed values of the data. The sample sizes are indicated in parentheses.

### 3-3. Effects of prolonged hypergravity exposure on the content of elements in the shoot system

Due to the observed alteration in root biomass levels under hypergravity conditions, there is a consideration that the uptake of mineral nutrients by roots might also be affected. We investigated the content of inorganic elements in the shoot by employing an ionomics approach using an ICP-OES instrument. Figure 6 shows effects of 40-d exposure to 10 *g* hypergravity on the distributions of inorganic elements in the main stems or the rosette leaves. As a result, no significant differences were found in either macronutrients (Ca, K, Mg, P), micronutrients (Cu, Fe, Mn, Mo, Zn), or harmful elements (Co) (Fig. 6a, b). Although not statistically significant, there was a slight tendency for Mo to increase under hypergravity conditions compared to the 1 *g* control (Fig. 6a, b).

**Fig. 6.**
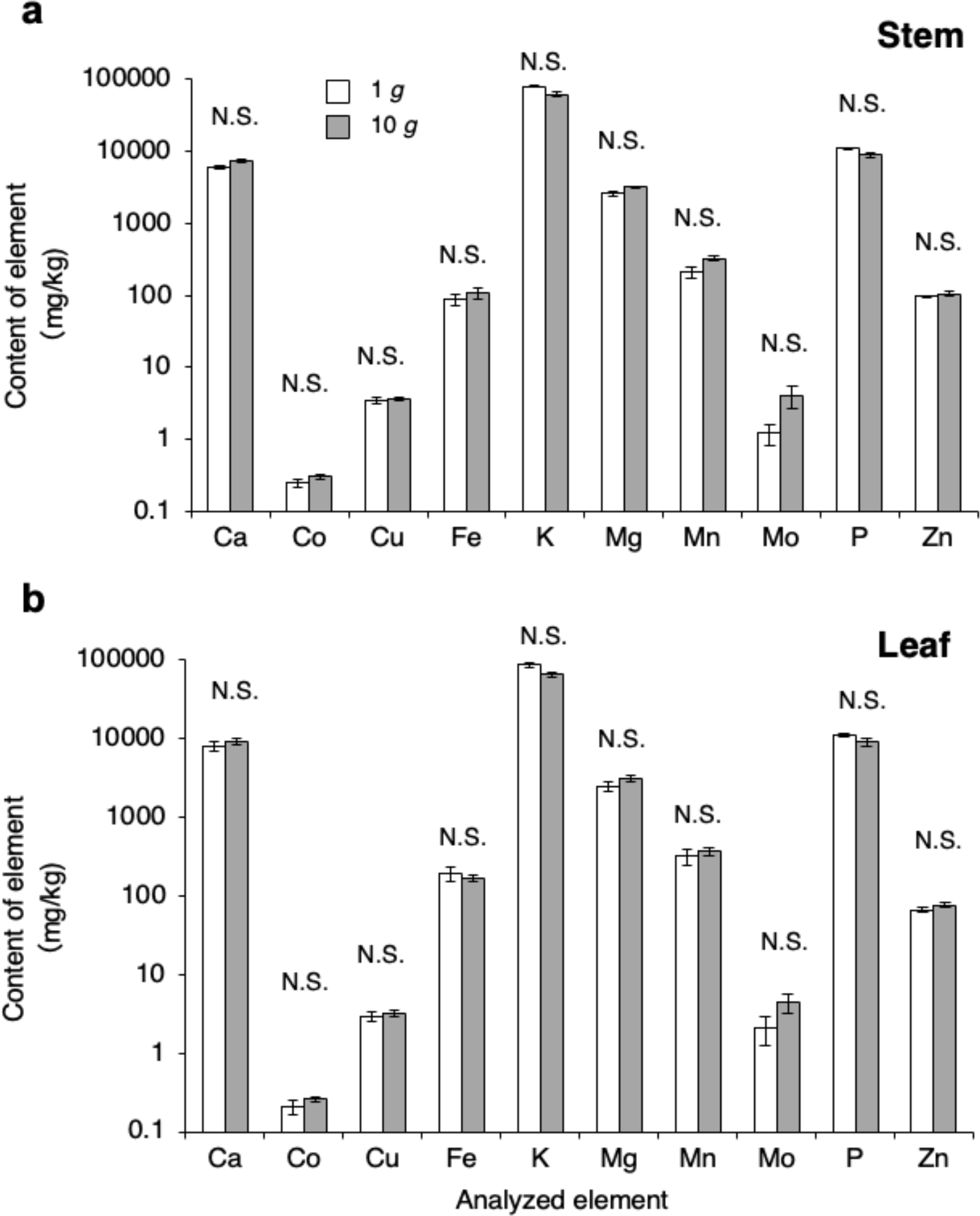
Effects of 40-d exposure to 10 *g* hypergravity on the content of elements (Ca, Co, Cu, Fe, K, Mg, Mn, Mo, P, Zn) in the shoot system of the Arabidopsis plants. White columns, 1 *g*; shaded columns, 10 *g*. Values are mean ± SE. N.S., *P* > 0.05, Wilcoxon rank sum test. (**a**) The main stem of the primary inflorescence. n = 4. (**b**) The rosette leaves. n=3 (1 *g*), 4 (10 *g*).

## 4. Discussion

We successfully cultivated Arabidopsis plants for an extended period of up to 40 d using an improved centrifuge designed specifically for prolonged hypergravity cultivation. The results showed that the length of the main stem of the primary inflorescence was significantly shorter in 40-d-old plants and tended to be shorter in 30-d-old plants compared to the 1 *g* control (Fig. 2a, Supplementary Fig. 1b). Additionally, the dry mass of the stem per unit length was significantly higher compared to the 1 *g* control for both cases (Fig. 2d, Supplementary Fig. 1d), which supports our previous findings obtained using Arabidopsis primary inflorescences treated with 300 *g* for 1 d (Nakabayashi et al., 2006; Tamaoki et al., 2006) and 8 *g* for 10 d (Shinohara et al., 2024). These effects are known as gravity resistance responses of the stem (Hoson and Soga, 2003; Soga, 2013). Additionally, the total dry mass of the stems of the lateral shoots was significantly higher under hypergravity compared to the 1 *g* control when examined using 30-d-old plants (Supplementary Fig. 1e).

While no significant difference was found in the number of rosette leaves between 1 *g* and 10 *g* conditions (Fig. 3b), the dry mass of the entirety of the rosette leaves was significantly higher under hypergravity compared to the 1 *g* control (Fig. 3c, Supplementary Fig. 1f). This indicates that the dry mass of each rosette leaf increased under hypergravity conditions. However, this increase is not solely attributed to the larger area of the rosette leaf under hypergravity compared to the 1 *g* control (Fig. 3d), but also to the significantly higher LMA of the rosette leaf under hypergravity compared to the 1 *g* control (Fig. 3g). From an ecophysiological standpoint, multispecies analyses demonstrate that elevated LMA is associated with increased intrinsic water use efficiency (Medrano et al., 2009; Gago et al., 2014). Consequently, the observed increase in LMA under hypergravity conditions, as compared to the 1 *g* control (Fig. 3g) implies an enhancement in water use efficiency under hypergravity.

A positive relationship is observed between photosynthetic capacity and LMA (Poorter et al., 2009). It has been reported that the area-based photosynthesis rate was enhanced under 10 *g* in the moss *P. patens* (Takemura et al., 2017; Takemura et al., 2017). Therefore, it is suggested that the photosynthesis rate is enhanced also in Arabidopsis, resulting in increased biomass of an individual plant (Fig. 5a, Supplementary Fig. 2a). It is also possible that enhanced photosynthesis rate led to increased rates of cell division and/or elongation under hypergravity. In a study involving cotyledons of *Ocimum basilicum* L., no significant effect on area and fresh mass was observed when exposed to 100 *g* for 2 d (Watanabe et al., 2024). Conversely, in our previous study (Shinohara et al., 2024), we observed an increase in lignin content in the rosette leaves of Arabidopsis when exposed to hypergravity at 8 *g* for 10 d during their development, in addition to the hypergravity effects observed in the present study. The difference between their findings and ours could be attributed to disparities in the duration of exposure or possibly variations in the sensitivity of plant species to gravitational acceleration.

In terms of biomass allocation, the Root-Shoot ratio was significantly higher under hypergravity conditions compared to the 1 *g* control (Fig. 5b, Supplementary Fig. 2b), and RMF was significantly higher under hypergravity conditions compared to the 1 *g* control in both 30-d-old and 40-d-old plants (Fig. 5e, Supplementary Fig. 2e). In *P. patens*, 83 % of the total biomass was allocated to shoots under 10 *g*, while 93 % was allocated to shoots under 1 *g* (Takemura et al., 2017), suggesting a relatively lighter shoot mass under hypergravity. Although this response contrasts with the findings in Arabidopsis revealed in the present study, this response of *P. patens* can also be interpreted as a potential adaptation to hypergravity, as reduced shoot mass might mitigate the risk of buckling under its weight (Kume et al., 2021). Conversely, the root system plays a crucial role in anchoring and supporting the shoot system. Therefore, the significant increase in the dry mass of the entire root system under hypergravity conditions indicates an enhancement of this anchoring and supporting function, contributing to gravity resistance mechanisms. The observed increase in Arabidopsis plant biomass at the individual level under hypergravity conditions (Fig. 5a, Supplementary Fig. 2a) suggests the potential for decreased biomass under low-gravity conditions, underscoring the need for countermeasures in plant cultivation on the moon or Mars.

Due to the evidence that the dry mass of the entire root system was significantly higher under hypergravity conditions compared to the 1 *g* control (Fig. 4b, Supplementary Fig. 1h), there was a consideration that the uptake of mineral nutrients by roots might also be affected. Interestingly, however, no significant differences were found in the levels of any of the elements analyzed (Fig. 6a, b). This suggests that homeostasis is maintained even under altered gravitational conditions. The finding that uptake of harmful elements such as Co is not necessarily enhanced is crucial from the perspective of plant cultivation using regolith on the moon or Mars.

## 5. Conclusions

We have successfully demonstrated that prolonged exposure to hypergravity for up to 40 days increases the biomass of Arabidopsis plants (the stem, rosette leaves, and roots) using a custom-built centrifuge equipped with a lighting system tailored for hypergravity cultivation. Specifically, we observed a significant increase in the dry mass of the stem per unit length under hypergravity, indicative of a typical gravity resistance response in the stem. Additionally, LMA of the rosette leaves significantly increased under hypergravity, suggesting an enhancement in photosynthesis rate and subsequent biomass increase in individual plants.

In terms of biomass allocation, both the Root-Shoot ratio and the RMF significantly increased under hypergravity. Furthermore, intriguingly, despite the increased dry mass of the root system, we found no significant differences in the content of any of the mineral elements analyzed in the shoot system between 10 *g* and 1 *g* conditions. This suggests that homeostasis of mineral nutrient uptake is maintained even under hypergravity conditions. These findings provide valuable insights for the design of plant cultivation systems aimed at future long-term manned space exploration missions to the moon or Mars.

## CRediT Author contributions

**Kazuki Ohara** and **Mizuki Katayama**: Investigation, Formal analysis, Validation. **Hiroyuki Kamachi**: Investigation and Data curation (ICP-OES), Writing – review & editing. **Atsushi Kume**: Methodology (Hypergravity cultivation system designing), Writing – review & editing. **Ichirou Karahara**: Conceptualization, Methodology (Hypergravity cultivation system designing), Writing – original draft, Writing – review & editing, Visualization, Funding acquisition, Supervision.

## Acknowledgements

This work was supported partly by JSPS KAKENHI Grant Nos. 21570064, 24K09514 (to I. K., H. K.), 23K17399 (to A. K., H. K., I. K.) and by Research Funding Grant by the president of University of Toyama. This work was supported by the Division of Instrumental Analysis at University of Toyama.

## Supplementary Materials

**Supplementary Fig. 1.**
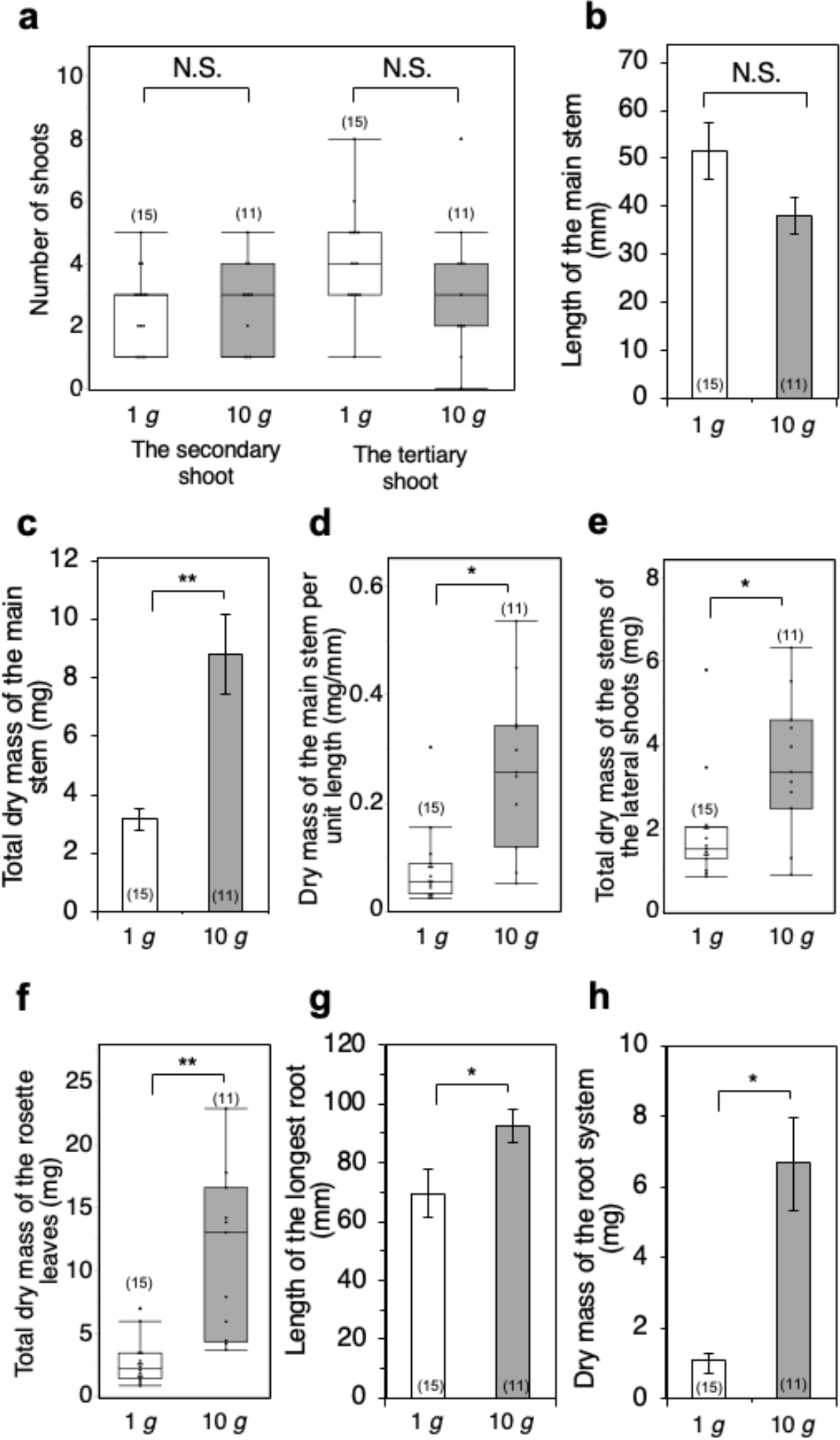
Effects of 30-d exposure to 10 *g* hypergravity on the morphology and biomass of the Arabidopsis plants. (**a**) The numbers of the secondary shoots and the tertiary shoots. N.S., *P* > 0.05, Wilcoxon rank sum test. (**b**) The length of the main stem of the primary inflorescence. N.S., *P* > 0.05, Welch’s *t*-test. (**c**) The total dry mass of the main stem of the primary inflorescence, **, *P* < 0.01, Welch’s *t*-test. (**d**) The dry mass of the main stem per unit length. *, *P* < 0.05, Wilcoxon rank sum test. (**e**) The total dry mass of the stems of the lateral shoots. *, *P* < 0.05, Wilcoxon rank sum test. (**f**) The total dry mass of the rosette leaves. **, *P* < 0.01, Wilcoxon rank sum test. (**g**) The length of the longest root. *, *P* < 0.05, Welch’s *t*-test. (**h**) The dry mass of the entirety of the root system. **, *P* < 0.01, Welch’s *t*-test. (**b**, **c**, **g**, **h**) Values are presented as mean ± SE.

**Supplementary Fig. 2.**
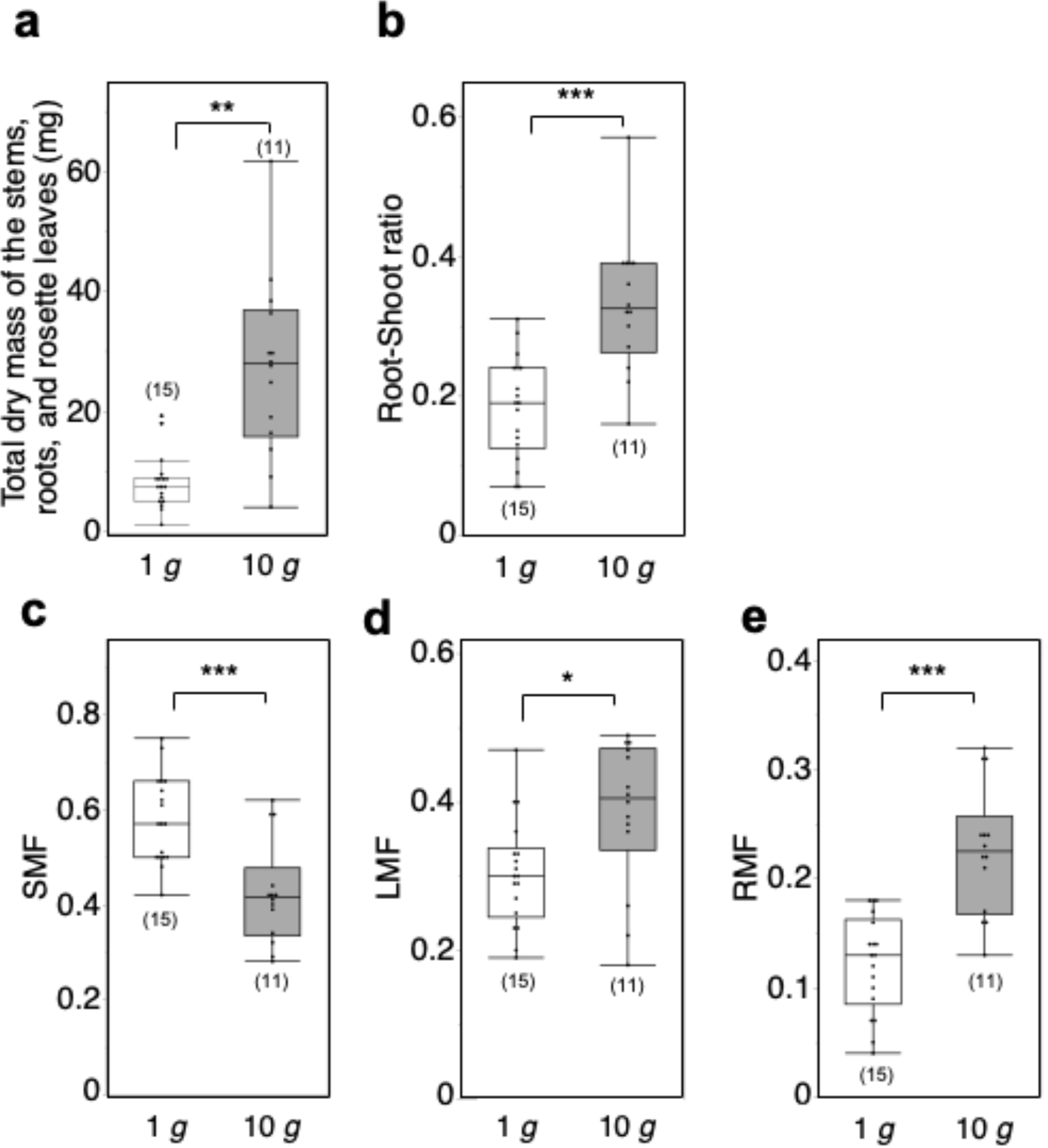
Effects of 30-d exposure to 10 *g* hypergravity on the biomass allocation of the Arabidopsis plants. (**a**) The total dry mass of the stems, roots, and rosette leaves. **; *P* < 0.01, Wilcoxon rank sum test. (**b**) Root-Shoot ratio. ***, *P* < 0.001, Student’s *t*-test of angler transformed values of the data. (**c**) SMF. ***, *P* < 0.001, Student’s *t*-test of angler transformed values of the data. (**d**) LMF. *, *P* < 0.05, Student’s *t*-test of angler transformed values of the data. (**e**) RMF. ***, *P* < 0.001, Student’s *t*-test of angler transformed values of the data. The sample sizes are indicated in parentheses.

